# Hiding in Plain Sight: Phylogenomics Reveals a New Branch on the Noctuoidea Tree of Life

**DOI:** 10.1101/2023.03.10.529269

**Authors:** Ryan A. St Laurent, Paul Z. Goldstein, Scott E. Miller, Robert K. Robbins

## Abstract

We present the results of the first phylogenomic analyses based on anchored hybrid enrichment (AHE) data from densely sampled tribes and subfamilies of Notodontidae (Prominent Moths). Our analyses reveal the family’s polyphyly with respect to an assemblage of genera related to *Scrancia* Holland that has been variously recognized at the tribal or subfamilial rank. We propose and re-describe Scranciidae, **stat. nov.**, and recognize 21 genera and approximately 100 species—distributed in Africa, Asia, and Australia and not represented in previous phylogenomic studies—from the six recognized noctuoid families (Noctuidae, Erebidae, Euteliidae, Nolidae, Notodontidae, and Oenosandridae). We further re-interpret morphological synapomorphies previously proposed for Notodontidae (including Scranciidae) and for the trifid Noctuoidea, *viz.* the ventral-facing tympanum and trifid forewing venation— characters previously called into question when Doidae were transferred from Noctuoidea to Drepanoidea. Deep-level relationships within Noctuoidea are not firmly established outside the clade comprising the four quadrifid families (Noctuidae, Erebidae, Euteliidae, and Nolidae), and in attempting to establish the phylogenetic position of Scranciidae relative to Notodontidae, Oenosandridae, and the quadrifids, we obtain conflicting results depending on data type (amino acid vs. nucleotide) and analytical framework (maximum likelihood, multi-species coalescent, and parsimony). We also demonstrate that discordant topologies among these ancient lineages yield drastically different divergence time estimates, highlighting the need for caution when interpreting phylogenetic dating of uncertain topologies. Following multiple analyses of several datasets designed around the distribution of missing data, and an evaluation of strict support measures at the deepest nodes of the noctuoid tree, we provisionally conclude that this ambiguity is a function of character conflict amplified by missing data and short branch lengths, and that in the topology best supported by the available data, Scranciidae is placed well outside Notodontidae and sister to the remaining Noctuoidea.

---

With over 40,000 described species (Kitching and Rawlins 1998; van Nieukerken et al. 2011), Noctuoidea (cutworms, loopers, owlets, prominents, tiger moths, and tussock moths) are the most species-rich, behaviorally diverse, and economically significant superfamily of Lepidoptera (butterflies and moths). Their economic importance is due to a high concentration of pests, such as the Old World Bollworm, *Helicoverpa armigera* Hardwick, the Corn Earworm, *Helicoverpa zea* (Boddie), the Spongy Moth, *Lymantria dispar* (L.), and the Fall Armyworm, *Spodoptera frugiperda* (J.E. Smith), the last of which recently colonized four continents in under five years (Tepa-Yotto et al. 2021). Although the composition of Noctuoidea has received considerable attention in molecular phylogenetic studies (Zahiri et al. 2011; Kobayashi and Nonaka 2016; Regier et al. 2017; Keegan et al. 2021), relationships among the six currently recognized noctuoid families—Erebidae, Euteliidae, Noctuidae, Nolidae, Notodontidae, and Oenosandridae—are not firmly established. Phylogenomic data have only recently begun to impact our understanding of the noctuoid backbone topology, but taxon sampling has been limited, considering the richness and diversity of this superfamily (Kawahara et al. 2019; Rota et al. 2022).

Historically, Noctuoidea classification was based largely on character systems associated with wing venation, tympanal configuration (an auditory structure), and larval morphology. The first of these is the eponymous character used to organize the noctuoid families into the “trifids,” in which forewing vein M_2_ originates from the discal cell midway between M_1_ and M3, and the “quadrifids,” in which vein M_2_ originates near the base of M_3_. Despite well-established groupings, phylogenetic progress has driven significant re-casting of both major groups. For quadrifids, the former Noctuidae were cleaved into four families, while subsuming Arctiidae and Lymantriidae as subfamilies within one of these (Erebidae). The established trifid classification recognized Notodontidae to include the previously family-level taxa Dioptidae and Thaumetopoeidae (Miller 1991). Although Doidae share trifid venation, tympanal configuration, and larval morphology with Notodontidae, this family was transferred to Drepanoidea based on molecular phylogenetic results (Bazinet et al. 2013; Kawahara et al. 2019; Rota et al. 2022). The small Australian family Oenosandridae was previously considered a subfamily of Notodontidae but has since been elevated to family rank (Miller 1991; Zahiri et al. 2011). Erebidae, Euteliidae, Noctuidae, and Nolidae comprise the quadrifids while Notodontidae and Oenosandridae represent the trifid lineages. Molecular phylogenetics have been brought to bear on all six currently recognized noctuoid families to varying degrees, including the two largest quadrifid families Erebidae and Noctuidae (Zahiri et al. 2013, 2023; Wang et al. 2015; Homziak et al. 2019; Keegan et al. 2019, 2021; Dowdy et al. 2020). The large family Notodontidae (4,500+ species), however, has received considerably less attention than the other large families, and notodontid monophyly has not been tested with phylogenomic data.

Research currently underway to develop a phylogenomic dataset for all major lineages of Notodontidae offers an opportunity to test classification hypotheses that have persisted since Miller’s (1991) phylogenetic analysis of adult and larval characters. In our efforts to sample exemplars of named (and suspected) subfamilial- and tribal-level groups in Notodontidae, we consistently obtained results placing the notodontid subfamily Scranciinae outside Notodontidae, and thus we recognize a need to revisit the character systems originally developed to support its placement. We formally raise the status of Scranciinae to Scranciidae, **stat. nov.**, reexamine Miller’s (1991) suites of characters supporting their former placement in Notodontidae, present a novel diagnosis and re-description of the group, and establish its composition.

Scranciidae, **stat. nov.**, are a strictly Old World family that includes roughly 100 species, found primarily in Africa and Southeast Asia. The adults are recognizable by their elongate legs and overall delicate form, and their larvae are especially noteworthy for their proleg structure that gives rise to a “looping” mode of locomotion similar to that observed in inchworms (Geometridae), loopers (Noctuidae), and some Erebidae (Holloway 1983). Scranciid larvae also have modified anal prolegs in the form of elongate stemapods, a trait frequently observed in Notodontidae. Originally flagged as an assemblage of atypical notodontid genera (“*Scrancia* group” *sensu* Gaede (1934), “*Gargetta* group” *sensu* Holloway et al. (2001)), the Scranciini were first formally assigned to Dudusinae (Miller 1991), and later elevated to subfamily rank by Schintlmeister (2008). Here we reassess some of the morphological evidence supporting a position for scranciids within Notodontidae and present novel data demonstrating that they are a distinct group displaying a combination of characters that require reinterpretation. We suggest that Scranciidae, **stat. nov.**, represent the sister group to all other Noctuoidea.

The removal of the scranciids from Notodontidae is significant in part because notodontids and scranciids share characters that have been considered nearly infallible synapomorphies. *Scrancia* Holland and its relatives share trifid forewing venation with Notodontidae and Oenosandridae, forming the “the trifid lineage” of Miller (1991), excluding Doidae. Scranciids also share a ventral-facing metathoracic tympanal membrane with Notodontidae (and again Doidae), whereas oenosandrids and the quadrifid families share an alternate state, with the tympana directed rearward. Interpretation of the distribution and transformations of these characters is central to our understanding of noctuoid evolution. For example, moth tympana are used to hear and are thought to play a major role in bat predator detection and avoidance (Kristensen 1998, 2003). Thus, the establishment of Scranciidae, **stat. nov.** and its phylogenetic placement presents several new challenges, upends the systematics of the group, forces reconsideration of character optimizations and evolution of key adaptations, and warrants caution when estimating divergence timing within the superfamily.

Far from stabilizing the already problematic backbone topology of the superfamily (Regier et al. 2017); recognition of Scranciidae appears to contribute novel sources of conflict, posing many issues for downstream studies. Using anchored hybrid enrichment phylogenetics (Lemmon et al. 2012; Breinholt et al. 2018), our goal is to illuminate an overlooked and understudied group, and one of the most species-rich Macroheteroceran families to be newly recognized in over a century. We test the hypothesis that Scranciidae does not belong within Notodontidae, as it was formerly classified; explore possible positions along the noctuoid backbone using a variety of methods investigating site, gene, and topological discordance; and test whether alternative topologies of the Noctuoidea backbone impact attempts to estimate divergence timing of this globally distributed, vast clade of insects.

## MATERIALS AND METHODS

### Taxon sampling

This project uses 52 Notodontidae, five Scranciidae, 15 species from other noctuoid families, and another 14 from six other superfamilies as outgroups (Table S1). Sequences were either previously published (Kawahara et al. 2019, St Laurent et al., in press), otherwise publicly available (Wellcome Sanger Institute 2022), or sequenced *de novo* from material in the McGuire Center for Lepidoptera and Biodiversity, Florida Museum of Natural History, University of Florida, Gainesville, FL, U.S.A. (MGCL); the Natural History Museum, London, UK (NHMUK); and the National Museum of Natural History, Smithsonian Institution, Washington D.C., U.S.A. (USNM). Scranciidae sampling includes taxa from across Africa and Asia, the two genera originally used to define the tribe Scranciini (*Scrancia* and *Gargetta* Walker), and other putative Scranciidae genera (all of these from the USNM). To help contextualize the placement of Scranciidae and test the monophyly of Notodontidae, the 45 notodontid samples cover all named subfamilies and tribes *sensu* Miller et al. (2018) using type genera whenever possible. We also include additional focused sampling in the notodontid subfamily Spataliinae (Ceirinae in Miller et al. (2018)), including type genus *Spatalia* Hübner, since that subfamily contains several genera erroneously treated as “Scranciinae” by previous authors.

### Morphology

Scranciidae identification follows Schintlmeister (2008, 2020) and Kiriakoff (1964). Much of the African Scranciidae have not been recently revised, and in some cases, specimens are identified only to genus. Differentiating this group and establishing its position outside Notodontidae requires revisiting the synapomorphies in Miller’s (1991) classification of the trifid Noctuoidea. Our approach, and our diagnosis of Scranciidae, **stat. nov.** draws from that work, supplemented with additional observations of specimens and images. Some of our terminology is based on interpretations of genitalia structures of Scranciidae in Schintlmeister (2008, 2020; see also File S1 and Table S2).

### DNA extraction, sequencing, and dataset preparation

Our pipeline for museum specimen DNA extraction, sample processing, sequencing, and downstream bioinformatics follows the notodontid study of St Laurent et al. (in press). The extraction protocol follows Hamilton et al. (2019), and the Lepidoptera-specific anchored hybrid enrichment (AHE) probe set used here follows Breinholt et al. (2018). All data, whether originally derived from specimens sequenced here or previously for AHE, or as publicly available genomes/transcriptomes, are combined with the AHE LEP1 loci, by using python scripts from Breinholt et al. (2018) to extract relevant loci from other published sources. The LEP1 kit itself consists of up to 855 highly conserved nuclear loci.

To test the monophyly of the Scranciidae and explore support for alternative placements within the Noctuoidea, we built 11 datasets with varied coverage and missing data, phylogenetic informativeness, and guanine-cytosine (GC) content. Importantly, in all datasets Scranciidae had relatively low missing data overall due to highly successful AHE target capture for that family (average of 10% missing data across the datasets). The datasets and their identifiers are: (1) all_NA, all nucleotides of loci recovered from the LEP1 set across all taxa included herein (854 loci, 202,140 base pairs (bp), parsimony informative sites = 41.1%, average missing data per terminal = 26.8%); (2) all_AA, translated amino acid residues for those data; (3) 60_percent_recovery_NA, nucleotides of loci recovered from ≥60% of taxa (689 loci, 162,390 bp, parsimony informative sites = 40.9%, average missing data per terminal = 19.4%); (4) 60_percent_recovery_AA, the translated amino acid residues for those data; (5) 90_percent_recovery_NA, nucleotides of loci recovered from ≥90% of taxa (153 loci, 39,042 bp, parsimony informative sites = 38.1%, average missing data per terminal = 5.9%); (6) 90_percent_recovery_AA, translated amino acid residues for those data; (7) Oeno_limited_NA, nucleotides of loci with 100% recovery for *Discophlebia* Felder, our most data-limited representative of Oenosandridae, a family of uncertain placement in Noctuoidea (392 loci, 92,940 bp, parsimony informative sites = 40.9%, average missing data per terminal = 15.2%); (8) Oeno_limited_AA, translated amino acid residues for those data; (9) NSORT, the 600 most informative loci (minimized root-to-tip variance, level of saturation, average patristic distance; maximized proportion of variable sites, average bootstrap across gene trees) according to genesortR (Mongiardino Koch 2021) (see Dataset subsampling in File S1); (10) AASORT, translated amino acid residues for those data; (11) NGC, which is the 60_percent_recovery_NA dataset with loci that had over 60% GC content removed.

### Phylogenetics

Phylogenetic inference was conducted in maximum likelihood (ML), parsimony, and multi-species coalescent (MSC) frameworks. For all ML analyses we employed IQ-Tree 2 v. 2.1.3 (Minh et al. 2020b). For all 11 datasets, IQ-Tree was run 100 times independently using models and partitions selected by a single run of ModelFinder (the additional command “-r cluster 10” was used in codon partitioned datasets to improve computation time) in IQ-Tree using the “-m TESTNEWMERGEONLY” command for each dataset (Kalyaanamoorthy et al. 2017), see File S1 for additional partitioning information. For all tree inference IQ-Tree runs, we calculated 1000 Ultrafast Bootstraps (UFBoot), 1000 SH-aLRT replicates, and called the “bnni” command to alleviate issues of model violation inherent in UFBoot (Minh et al. 2013, 2020b; Hoang et al. 2018a). A separate IQ-Tree run was completed on the 60_percent_recovery_NA dataset partitioned and modeled as before for this dataset (initially by locus then partitions consolidated with ModelFinder), but with 100 standard nonparametric bootstraps instead of UFBoot and transfer bootstrap (TBE) supports (“--tbe” command) as further alternative measures of supports (Felsenstein 1985; Lemoine et al. 2018).

For topology tests (below and File S1) we evaluate topologies observed across analyses and determine whether any are preferred or that could be rejected *a priori*. Using IQ-Tree, we performed a separate set of analyses with 60_percent_recovery_NA and 60_percent_recovery_AA datasets partitioned as above, in each case constraints passed to IQ-Tree with the “-g” command. These constrained analyses forced the tree search to maintain the relative relationships of Scranciidae, Oenosandridae, Notodontidae, and quadrifids (rooted to non-noctuoid outgroups) as identified by various recovered topologies being tested.

Initial parsimony analyses of the 60_percent_recovery_NA and 60_percent_recovery_AA datasets were undertaken in MPBoot v. 1.1.1 (Hoang et al. 2018b) with 1000 MPBoot bootstrap replicates in calculated for support. These were followed by analyses in TNT v. 1.6 (Goloboff et al. 2008; Goloboff and Catalano 2016), which were used to compare branch lengths, Bremer support, and jackknife values on the 60_percent_recovery_NA dataset. Both traditional and New Technology searches (sectional, ratchet, drift, and tree-fusing) were undertaken in TNT.

Finally, as an alternative to concatenation methods, we inferred a species tree under the MSC in ASTRAL III v. 5.7.8 (Zhang et al. 2017). Individual gene trees were the same as those passed to genesortR, but for MSC analysis we used only the gene trees derived from the 60_percent_recovery_NA dataset. Support values in ASTRAL are posterior probability “Astral Support Values” (ASV).

In the present study we consider the following levels of support to be “robust” for each type of analysis: UFBoot ≥ 95 and SH-aLRT ≥ 80; ASV, MPBoot supports, nonparametric BS, and TBE ≥ 95, largely based on documentation of the respective programs.

### Concordance and topology tests

Due to multiple topologies of Noctuoidea obtained in previous studies (Mutanen et al. 2010; Zahiri et al. 2011; Regier et al. 2017; Kawahara et al. 2019; Keegan et al. 2021) and in the present work (see Results, Figure S1), we examined concordance between sites, genes, species trees, and gene trees as well as several topology tests to explore whether there is a basis for a preferred topology for downstream analyses.

To investigate locus- and site-specific support for nodes in our results, we calculated gene concordance factors (gCF) and site concordance factors (sCF) (Minh et al. 2020a). The latter were restricted to the best of 100 IQ-Trees inferred on the concatenated 60_percent_recovery_NA dataset partitioned according to locus, without merging of partitions. Gene trees for gCF and sCF were the same as those used for our genesortR and MSC analyses, but only those from the 60_percent_recovery_NA dataset. Finally, we performed discordance analyses on species trees and a relative frequency analysis in DiscoVista (Sayyari et al. 2018).

IQ-Tree also permits the statistical testing of likelihoods of user-provided topologies to identify if a constrained tree (e.g., a topology resulting from a tree search constrained to a competing topology) is not significantly worse than an unconstrained tree. We used the Approximately Unbiased test (p-AU) to identify which topologies could be rejected. While topology tests help to identify which, if any, topologies can be rejected, IQ-Tree also includes Four Cluster Likelihood Mapping (FCLM) (Strimmer and von Haeseler 1997; Minh et al. 2020b). This method depicts the distribution of the most informative quartets and was used to identify the clade in which Scranciidae most frequently falls. Our four clusters were defined as (1) outgroups, (2) Oenosandridae, (3) Scranciidae, and (4) quadrifids + Notodontidae. The sister grouping of quadrifids and Notodontidae is here deemed appropriate since it appears in most of our results, and previous molecular and morphological research has congealed around this relationship (Miller 1991; Zahiri et al. 2011; Regier et al. 2017; Kawahara et al. 2019).

### Divergence time estimation

Since we recovered multiple topologies for the backbone of Noctuoidea (see Results), we date several possible trees independently rather than picking a tree for dating analyses *a priori*. Our dating method follows the ML method employed by St Laurent et al. (2021), which includes a custom pipeline of python scripts for running multiple TreePL (Smith and O’Meara 2012) analyses in parallel to combine into a single “consensus” dated tree. This allows for ranges of divergence times at every node, which is useful since a single TreePL run obtains only a single age at each node (see File S1).

## RESULTS

### Differentiation and phylogenetic position of Scranciidae

All analyses support the monophyly of Noctuoidea, Scranciidae, and each of the remaining noctuoid families for which multiple exemplars were sequenced (only single exemplars were available for Nolidae and Euteliidae) as in previous studies, where relevant (Zahiri et al. 2011, 2011; Regier et al. 2017). The monophyly of Scranciidae is accompanied by maximum bootstrap support in all analyses, regardless of dataset and analysis parameters, with the placement of Scranciidae variable but always outside and rarely the immediate sister to Notodontidae (Fig. 1). Based on these results and following enumeration of characters, shared and unshared with Notodontidae (Table S2), we re-describe and present an expanded diagnosis of Scranciidae.

**FIGURE 1.**
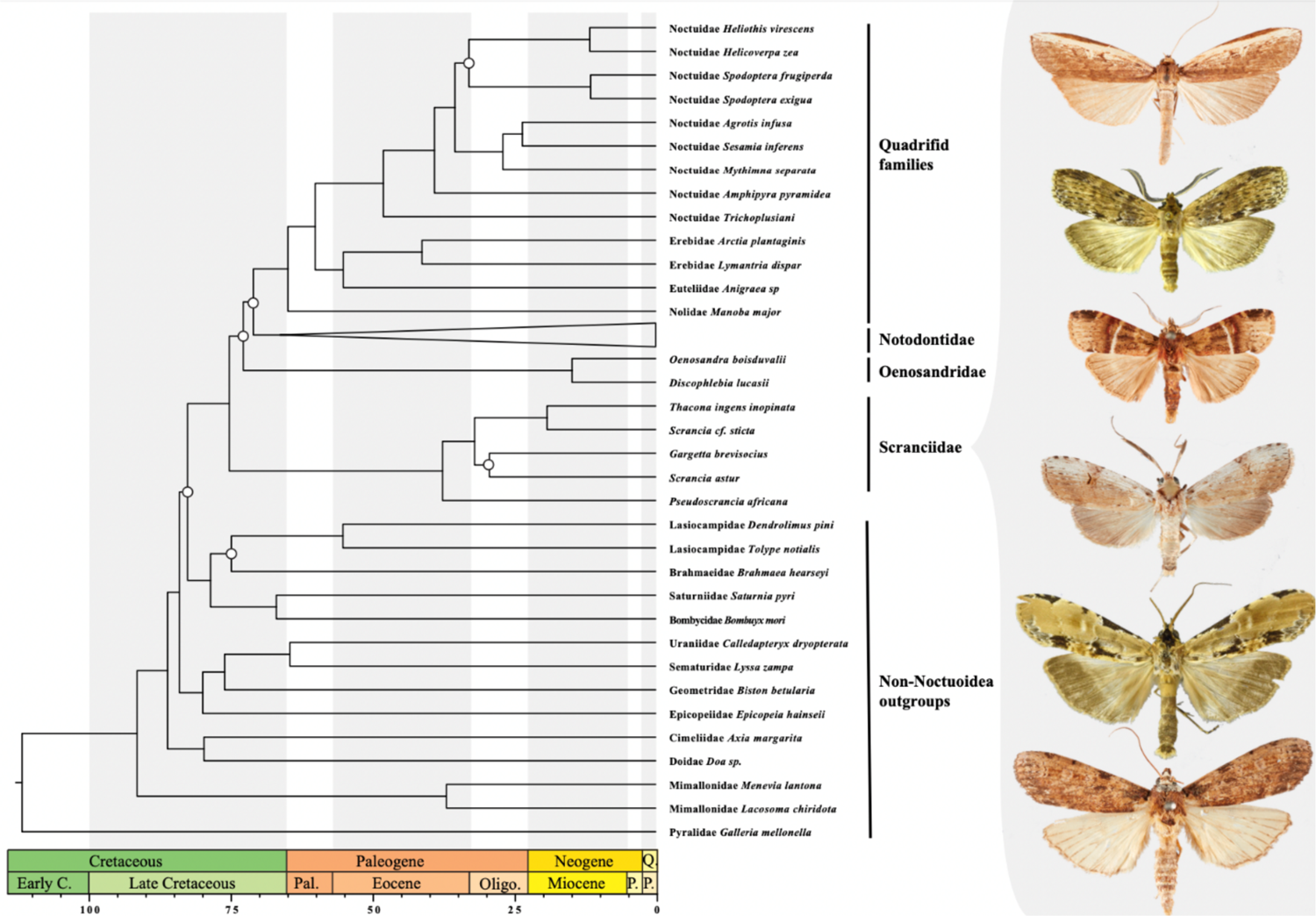
Chronogram of Noctuoidea and outgroups, phylogeny inferred in IQ-Tree on the 60_percent_recovery_NA dataset, dated with TreePL. Topology A is shown, with Scranciidae sister to all Noctuoidea. Representative Scranciidae genera to the right of the figure from top to bottom: *Phycidopsis*, *Gargetta*, *Gargettoscrancia*, *Lasioceros*, *Thacona*, and *Scrancia* (all in USNM, not to scale). For complete tip information, dates, and support values see Figures S2 and S3.

In evaluating alternative positions for the Scranciidae within the Noctuoidea, we focus on the four-taxon statement comprising the three trifid families (1) Scranciidae, (2) Oenosandridae, and (3) Notodontidae; and (4) the clade consisting of the four quadrifid families. While the monophyly of Noctuoidea and of each of these four component clades is well-established, the relationships among them are not. From our various analyses of both nucleotides or amino acid residues, under varying degrees of missing data, informativeness, and GC content, using different inference frameworks (ML, parsimony, or MSC), we recover seven topologies, A–G: (A) (Scranciidae, (Oenosandridae, (Notodontidae, quadrifids))) (shown in Fig. 1); (B) (Oenosandridae, (Scranciidae, (Notodontidae, quadrifids))); (C) (Oenosandridae, (Notodontidae, (Scranciidae, quadrifids))); (D) (Oenosandridae, (quadrifids,(Scranciidae, Notodontidae))); (E) ((Oenosandridae, Scranciidae), (Notodontidae, quadrifids)); (F) (Scranciidae, (Notodontidae, (Oenosandridae, quadrifids))); and (G) (Scranciidae, (quadrifids, (Notodontidae, Oenosandridae))). See Table S3 for an overview of the dataset type and inference method, Figure S1 for a diagram of competing topologies, and Dryad for tree files for topologies resulting from all analyses.

Only topology D supports a sister relationship of Scranciidae and Notodontidae, which is noteworthy given the recent inclusion of Scranciidae in Notodontidae. But bootstrap support for the MRCA of Notodontidae and Scranciidae in this topology is low in those analyses (Oeno_limited_AA: UFBoot/SH-aLRT: 54.9/62 and all_AA: 15.3/44). When the dataset is substantially reduced to 90_percent_recovery_AA, we recover Scranciidae as sister to the notodontid subfamily Platychasmatinae, thereby rendering Notodontidae paraphyletic with low support (63.9/57). This is the only case where Scranciidae are nested within Notodontidae, but we caution that 90_percent_recovery_AA is our smallest dataset, and these results may be spurious due to artificially limiting the dataset.

Regarding the most consistently recovered topology (A), in all cases we obtained high bootstrap values for the branches subtending Scranciidae and their immediate sister and Oenosandridae and their immediate sister. Robust bootstrap values are seen for this topology under UFBoot, SH-aLRT, standard nonparametric bootstraps, and transfer bootstraps (Table S3). Among ML analyses, this is the only topology where both roots appear supported simultaneously; in all other topologies the bootstrap values are highly skewed towards one or the other.

In parsimony analyses in TNT, branch lengths and support values for the two nodes subtending the primary divisions within Noctuoidea were markedly lower than those immediately above and below them: Branch lengths 859 and 865 relative to 3103/2732 and 1369/897 above, 1213 below; Bremer values 81 and 78 relative to 876/1000+ and 170/331 above, 437 below; and jackknife values 69 and 72 relative to 100/100 and 98/93 above, 100 below.

In view of possible discordance among data within Noctuoidea, we calculated concordance factors for the 60_percent_recovery_NA dataset that gives rise to topology A for the purpose of identifying whether sites or genes could be responsible for observed discordance across analyses. For that analysis we identified lower levels of gCF for two ancestral nodes of Noctuoidea (2.47 for the MRCA of superfamily and 0.842 for the MRCA of all Noctuoidea excluding Scranciidae) than of sCF (37 and 34.9, respectively), suggesting wider conflicts among loci than among individual sites. Locus-level discordance is explored more below.

Topology tests were carried out in IQ-Tree with the 60% recovery datasets (nucleotide and amino acids) in which the unconstrained topology (A) was tested against constrained topologies (B–G). The purpose of these analyses was to identify whether the statistical support for topology A was higher than for any competing constrained topology. Datasets 60_percent_recovery_NA and 60_percent_recovery_AA were tested separately to allow comparison of likelihoods. With nucleotides, only topologies A and G (both of which have Scranciidae sister to all other Noctuoidea) are not significantly excluded (p-AU for A = 0.919, B = 0.0201, C = 0.0169, D = 0.000487, E = 0.0185, F = 0.0275, G = 0.222). With amino acid residues, none of the competing topologies were significantly excluded (p-AU for A = 0.737 B = 0.347, C = 0.284, D = 0.539, E = 0.201, F = 0.313, G = 0.385). Four cluster likelihood mapping strongly supports (69.4%) the unrooted quartet of (Scranciidae,Outgroups) | (Oenosandridae,(quadrifids,Notodontidae)), which corresponds to topology A if rooted to the outgroups.

Our analyses with DiscoVista served three purposes: (1) to visualize discordance across analyses, (2) to further evaluate quantitative evidence for locus-level discordance, and (3) to illustrate the distribution of GC content across taxa since high GC content has shown to lead to topological discordance in Lepidoptera phylogenetics (Rota et al. 2022). For the first point, in ML and parsimony analyses, high bootstrap values are obtained for the monophyly of Oenosandridae, quadrifids, Scranciidae, and Notodontidae, although bootstrap support for Notodontidae is weaker when datasets are reduced (90_percent_recovery_AA and Oeno_limited_AA), at which point support also breaks down considerably with respect to the various possible placements of Scranciidae and Oenosandridae across Noctuoidea. Our second DiscoVista analysis shows the relative frequency of quartets that support a given topology with respect to a focal branch in the species tree (DiscoVista outputs available on Dryad). A majority of quartets generated from the 60_percent_recovery_NA loci support the relationship at branch number 6 (here denoted by a pipe, see DiscoVista figure on Dryad) that is observed in topology A: (Outgroups, Scranciidae) | (Oenosandridae, (quadrifids, Notodontidae)) but a majority of quartets also support the following arrangement with respect to branch number 7: (Notodontidae, Scranciidae) | (remaining Noctuoidea). If all_NA gene trees are examined with DiscoVista, the same result is observed for branch 6, but for branch 7 there are roughly equal frequencies of two alternative topologies. Finally, DiscoVista allowed us to identify that the focal taxa (Scranciidae) had among the lowest GC content of all taxa studied here, and therefore we do not consider that high GC content being the reason for discordance. Furthermore, the analysis carried out on the NGC dataset further supports this as removal of high GC loci from the dataset does not change the topology (topology A still observed).

### Dating

Table S3 displays the divergence times of key nodes from the various TreePL analyses, which were conducted for topologies A–C, to illustrate how discordant topologies can impact divergence time estimates. Importantly, we observed older crown and stem ages for the four focal clades in the single amino acid dating analysis than in the two nucleotide-based analyses. In all analyses, Noctuoidea crown ages were in the Cretaceous, with topology only minimally affecting mean ages. Specific internal nodes of Noctuoidea varied more noticeably, particularly for the long-branched Scranciidae. The crown age for the newly recognized family was recovered in the late Eocene-early Oligocene, with means ranging from the younger 33.4 Ma (topology C) to older ages 37.5 Ma and 39.8 Ma (topology A and B respectively) (dated topology A is shown in Fig. 1). The most pronounced difference among analyses was found in the stem age of Scranciidae, which ranged from 66.5 Ma in topology C to ∼75 Ma in topologies A and B.

### Morphological corroboration

Differentiating Scranciidae from Notodontidae requires reexamining the distributions and coding of characters adduced in support of alternative topologies uniting them with dudusine notodontids (Miller 1991), and assessing the implications of a novel topology for historically regarded synapomorphies of both trifid and quadrifid Noctuoidea *sensu* Kitching and Rawlins (1998). Based on these results and following enumeration of salient morphological characters presented by Miller (1991), we present a revised diagnosis for Scranciidae, **stat. nov.** below. The redescription is presented in File S1, and characters bearing on the monophyly of Scranciidae and related groups are summarized in Table S2.

**Scranciidae (Miller, 1991), stat. nov.**

Scranciini Miller, 1991: 185

Scranciinae Miller, 1991, in Schintlmeister 2008:8

Scrantiinae Miller, 1991, in Kobayashi 2021: 124

Type genus: *Scrancia* Holland, 1893

### Diagnosis

Adults Scranciid moths are most easily recognizable by their narrow forewings with straight or convex margins, large sub-triangular hindwings, overall thin and delicate build, extremely long and thin legs, broad accessory cell on the forewing (Fig. S4a–d), protruding frons (Fig. S4e–g), and male terminalia that (Fig. S4i–l), unlike other trifid noctuoids, often lack socii. While Scranciidae and Notodontidae (and drepanoid Doidae) share a ventral-facing tympanum and the absence of nodular sclerite previously thought to have been synapomorphies of Notodontidae, and trifid wing venation (considered here symplesiomorphic in Noctuoidea), the combination of the elongate, broad, accessory cell, absence of cteniophores, and frequent absence of socii is diagnostic of Scranciidae. The corpus bursae of many scranciid females bears a coffee bean-shaped signum (Fig. S4h) apparently unique to scranciids. Adult Scranciidae are differentiated from Old World Notodontidae and Oenosandridae by the “upright” posture *in vivo* with the anterior part of body held at a roughly 45° angle to the substrate with the mesathoracic femora reaching the head when held appressed to the thorax (Fig. 2a–e).

**FIGURE 2.**
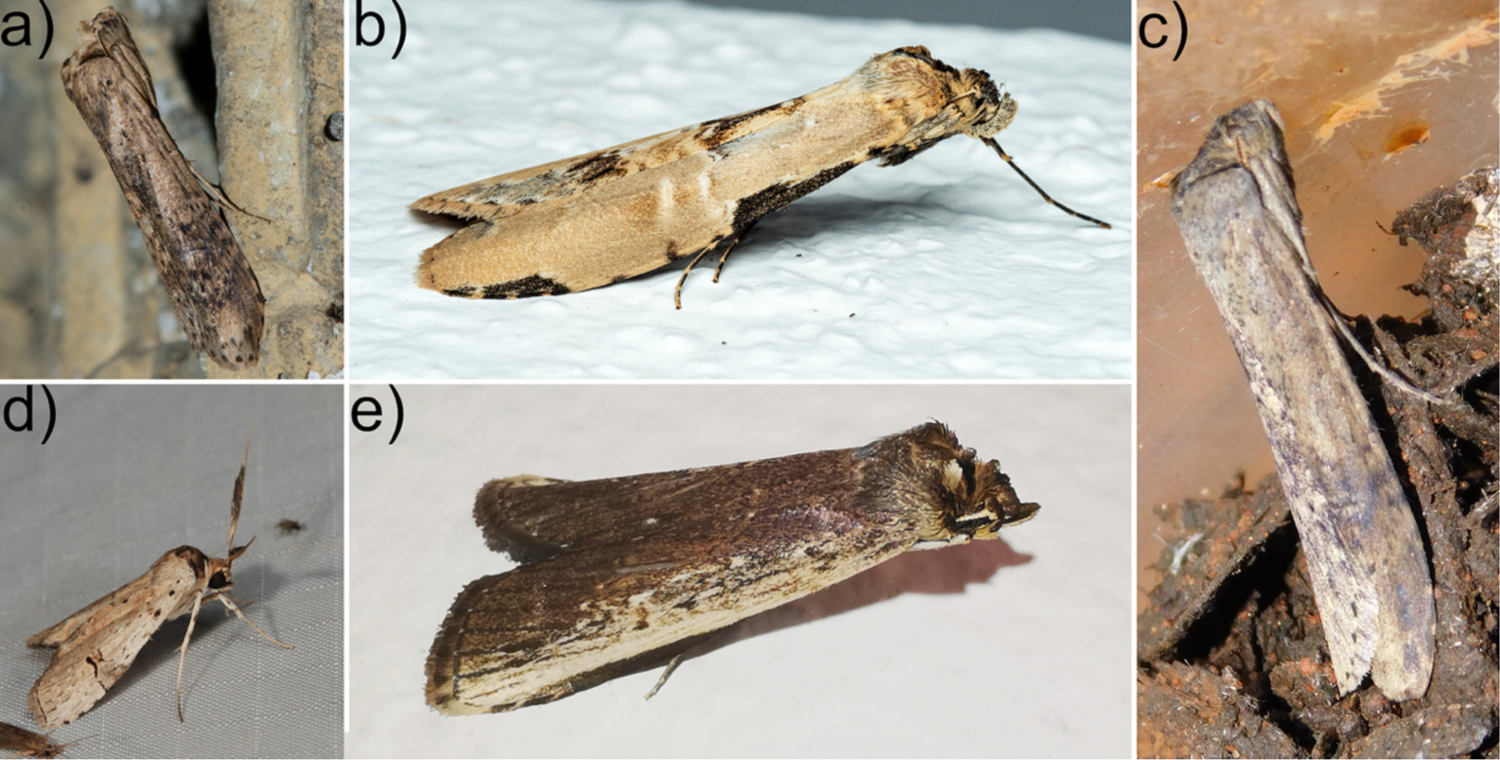
Adult Scranciidae *in situ*. a) *Gargetta sp.* (V. A. Ismavel). b) *Thacona ingens* (M. Sone). c) *Scrancia sticta* (Q. Grobler) d) *Lasioceros aroa* (D. Fischer). e) *Phycidopsis albovittata* (C. Stinchcomb).

Larvae. Scranciid caterpillars are unmistakable in having both reduced prolegs on abdominal segments three and four and stemapodiform anal prolegs with extendable filaments on the tenth abdominal segment (Fig. 3). The larvae are overall narrow and elongate with a comparatively large head, and only abdominal segments 4–6 or 5–6 being used for locomotion, first anchoring the larva while the thoracic segments with the true legs are thrust forward, and then “looped” upward as in inchworms (Geometroidea) (Holloway 1983). Within the Noctuoidea, the combination of reduced prolegs on A3 and A4 plus stemapods on A10 is unique to Scranciidae, excepting *Selenisa* and its close relatives in Erebidae: Ophiusini. However, the stemapods in Scranciidae are minutely spined or otherwise rugose, whereas those in the erebids are smooth (but may bear setae) and truncated.

**FIGURE 3.**
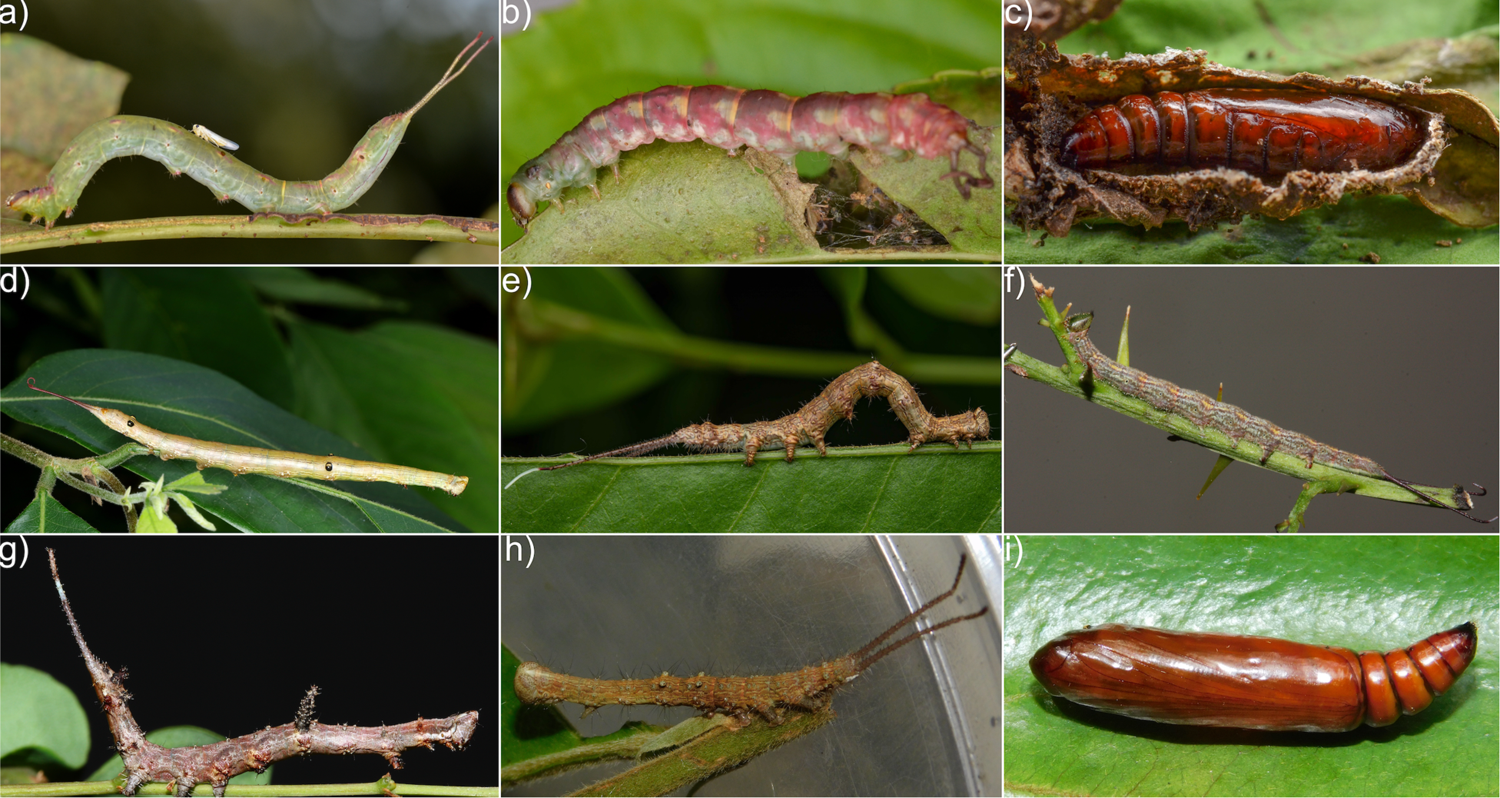
Immature stages of Scranciidae *in situ*. a) Larva of *Gargetta curvaria* (W. Jiajie). b) Prepupal larva of *G. curvaria* (W. Jiajie). c) Pupa of *G. curvaria* (W. Jiajie). d) Larva of *Gargetta cf. costigera* (SK LAI). e) Probable *Gargetta* (A. Tomaszek). f) Larva of *Phycitimorpha dasychira* (I. Sharp). g) *Postscrancia discomma* (I. Sharp). h.) *Scrancia sticta* (Q. Grobler). i) Pupa of *S. sticta* (Q. Grobler).

### Life history

The majority of larval food plant records are in the chemically defended, lactiferous Euphorbiaceae, but Anacardiaceae, Fabaceae, Ochnaceae, Phyllanthaceae, and Rutaceae are also eaten (Holloway 1983; Kroon 1999; Holloway et al. 2001; Schintlmeister and Witt 2015; Schintlmeister 2020); all of these plant families fall within either Fabales, Malpighiales or Sapindales (Chase et al. 2016).

### Distribution

The family is distributed in Africa south of the Sahara, India, Sri Lanka, Southeast Asia ranging into southern China, the Philippines, Indomalaya, and Australasia (Kiriakoff 1964; Holloway 1983; Schintlmeister 2008, 2020; Schintlmeister and Witt 2015). The group is mostly tropical.

### Composition of Scranciidae

Based strictly on our sequencing, we can conclusively include *Gargetta*, *Pseudoscrancia* Strand, 1915, *Scrancia*, **stat. rev.**, and *Thacona* Walker, 1865 in Scranciidae. Schintlmeister and Witt (2015) synonymized the African genus *Scrancia* (type genus of Scranciidae) with the Asian genus *Thacona* based on similarity in genitalia, but our analysis, which includes a single Asian *Thacona* and two African *Scrancia*, indicates that this renders Schintlmeister and Witt’s (2015) concept of *Thacona* paraphyletic (Fig. 1), and we therefore reinstate *Scrancia* as a valid genus with no Asian species. We caution that significantly more taxonomic study is needed, in our treatment, *Scrancia* still appears polyphyletic, and at least one new genus will likely be required to address this polyphyly.

Additionally, based on established apomorphies summarized here, we formally include within the Scranciidae 17 genera beyond those that were either treated by Miller (1991) or sequenced by us: *Archistilbia* Kiriakoff, 1954, *Dinotodonta* Holland, 1893 (Fig. S4d), *Gargettoscrancia* Strand, 1912 (Fig. 1), *Hijracona*, *Lamoriodes* Hampson, 1910, *Lasioceros* (Figs 1, 2d, S4g), *Leptonadatoides* Kiriakoff, 1969, *Lomela* Kiriakoff, 1962, *Malgadonta* Kiriakoff, 1962, *Noctuola* Schintlmeister, 2020, *Parascrancia* Kiriakoff, 1969, *Phycidopsis* (Figs 1, 2e), *Phycitimorpha* Janse, 1920 (Figs 3f, S4c), *Postscrancia* Schintlmeister and Witt, 2015 (Fig. 3g), *Stictogargetta* Kiriakoff, 1968, *Subscrancia* Gaede, 1928, and *Turnacoides* Gaede, 1928. Some, but not all, of these genera have been included in “Scranciinae” in recent works (Schintlmeister and Witt 2015; Schintlmeister 2020), notably Kobayashi and Nonaka (2016) recovered *Lasioceros* and *Phycidopsis* in their Scranciinae clade. Combined with the sequenced taxa, 21 genera are here assigned to Scranciidae. Table S4 summarizes these generic assignments. Genera formally assigned to Scranciini/Scranciinae but which we exclude based on phylogenomic results are discussed further in File S1.

## DISCUSSION

Phylogenomic data have resolved many relationships within Lepidoptera (Breinholt et al. 2018; Hamilton et al. 2019; Homziak et al. 2019; St Laurent et al. 2020; Rota et al. 2022). Revision of Noctuoidea relationships was initiated in the past decade on the basis of molecular phylogenetics (Zahiri et al. 2011, 2013, 2023; Keegan et al. 2021). We have not yet arrived at a stable resolution for Noctuoidea nor do we claim to here, given our sampling focus on Notodontidae and the inherent ambiguity in the available data, but the recognition of Scranciidae as a trifid lineage potentially independent of Notodontidae is warranted, and the possibility of its placement as sister to the remaining Noctuoidea is noteworthy. The elevation of Scranciidae represents a substantive rearrangement for the Noctuoidea, and the fifth intrafamilial group in the Noctuoidea to be elevated to family rank since 1991 following Erebidae, Euteliidae, Nolidae (all former subfamilies of Noctuidae), and Oenosandridae (formerly a belonging to “Thaumetopoeidae,” which themselves are now recognized as a notodontid subfamily). Scranciidae is the largest such group to have been removed from Notodontidae or be recognized within the trifids.

We have provided genetic and morphological evidence that these two groups should be treated as separate families, contradicting previous morphological and two-gene studies that maintained Scranciini/Scranciinae within Notodontidae (Miller 1991; Kobayashi and Nonaka 2016). Kobayashi and Nonaka (2016) employed limited outgroup sampling across Noctuoidea and only two genetic markers, warranting verification of the placement of Scranciinae. Our findings corroborate in part their results, which retrieved the scranciids distinct from Dudusinae and outside (but sister to) all remaining notodontids. They followed Schintlmeister’s (2008) elevation of Scranciinae from within Dudusinae. Based on our results, we instead propose the elevation to family, excluding Scranciidae from Notodontidae based on unshared morphological apomorphies (Table S2), and strong molecular support for a position even more distant from the Notodontidae. With the observation that scranciids depart significantly from established synapomorphies of the Notodontidae and that primary shared characters (putative synapomorphies) have elsewhere been demonstrated as homoplastic by virtue of the phylogenomic placement of Doidae outside of Noctuoidea, we conclude that support for the placement of scranciids within the notodontids was spurious. The characters shared by scranciids and notodontids are here deemed polymorphic (e.g., sclerotized pleuron of the eighth abdominal segment in females) and are interpret as either symplesiomorphies (presence of large ocelli and ventral-facing tympana) or secondarily derived (stemapods) (Miller 1991).

While the recognition of Scranciidae is straightforward, it is perhaps no surprise that substantial data do not offer an unambiguous solution to its placement. Based on our analyses, the placement of Scranciidae varies with the kind and quantity of data (amino acid vs. nucleotide, variously truncated to adjust percent coverage of loci, removal of high GC content loci, among other filtering steps), as well as inference method (ML, parsimony, MSC).

Scranciidae are sister to Notodontidae (ML analyses of three amino acid datasets), to Oenosandridae (parsimony analyses of nucleotides), to the quadrifid group (ML analysis of the most reduced nucleotide dataset, 90_percent_recovery_NA, and parsimony analysis of 60_percent_recovery_AA), to [Notodontidae + quadrifids] (ML analysis of 60_percent_recovery_AA), or finally, to all the other Noctuoidea (ML analyses of the most complete nucleotide datasets and by ASTRAL). Importantly, we note that only the three ML amino acid datasets that were most data-limited support their placement as sister to the Notodontidae, whereas under ML, all but the most reduced (90_percent_recovery_NA) nucleotide datasets support the placement of Scranciidae as sister to all remaining Noctuoidea, as does ASTRAL. Under no analytical circumstances did our analyses recover monophyly of the trifids, whereas quadrifids are consistently monophyletic.

Character signal/conflict, base pair composition, and model violation may all contribute to a given dataset’s lack of decisiveness with respect to a particular topology or node. Relatively low Bremer and jackknife values imply strong character conflict, no doubt amplified by short branch lengths. Although these values refer to nodes recovered only in parsimony (topology E), there is obviously even less character support for alternative (longer) trees, and branch lengths for nodes subtending alternative placements of Scranciidae are even shorter. Our fundamental interpretation of the different analytical results is that the impacts of character conflict and multiple short branches subtending very old taxa may be compounded by imperfect locus-level coverage that is characteristic of target capture data (hence the need to explore locus-level coverage). We observe, first, that results tend to vary more across inference methods than datasets, but that the matrices with the most data (or the best data, according to genesortR) converge on topology A under ML and on topology E under parsimony, and that these solutions differ only in the placement of Oenosandridae, where the recovery of Scranciidae apart from Notodontidae is consistent. We see a particularly strong analytical impact of nucleotides versus amino acids, particularly as the number of loci increases or when better quality loci are used (NSORT dataset determined by genesortR). However, for amino acids, our topology tests in IQ-Tree find all seven topologies equally likely since none can be reasonably excluded. Thus, nucleotides may be more decisive over much of the Noctuoidea backbone topology and result in less topological instability, especially with respect to the placement of Scranciidae. Synonymous change and base saturation may be playing roles considering the multiple short branches near the noctuoid base and the especially long branch represented by Scranciidae. This is further exemplified by the marked differences observed topologically in the Noctuoidea when different datasets are employed. The NSORT dataset minimizes saturation and still results in topology A with a subsampled set of 600 loci. The short branches along the noctuoid backbone may also reflect rapid radiation during the late Cretaceous, but evaluating this requires denser sampling.

Analyses in DiscoVista may suggest a possible explanation for the high degree of discordance across analyses in the present study. Our relative frequency analysis indicates that the frequency of quartets around key internal branches of topology A (those being the stem subtending all noctuoids except Scranciidae and the stem subtending [quadrifids + Notodontidae]) are each supported by a higher than null (1/3) frequency of quartets, but in neither case with overwhelmingly high frequencies. We also observe similar frequencies in the other two possible quartets on either branch, such an observation may be indicative of incomplete lineage sorting assuming gene tree error is not a major factor (Sayyari et al. 2018).

Given the branch lengths, apparent character conflict implied by gCF, DiscoVista, and multiple support measures, the data do not support an unambiguous placement of Scranciidae. Although we cannot rule out their placement as sister to Notodontidae, the paucity of unreversed morphological synapomorphies uniting the two justify the decision to erect a new family regardless of its position. More comprehensive sampling across Noctuoidea, and especially across Scranciidae and Oenosandridae, perhaps with full genomes, will be needed to provide further evidence to support a preferred topology. However, we hasten to register our preference for topologies A or G, in which Scranciidae are sister to the other Noctuoidea since they accommodate multiple analyses with the most data (60_percent_recovery_NA codon or locus partitioned, Oeno_limited_NA, NSORT, all_NA), cannot be statistically rejected in nucleotide topology tests, and have the highest observed combined UFBoot and SH-aLRT for both the placement of Scranciidae and Oenosandridae, the two most troublesome clades (Table S3). Furthermore, our FCLM overwhelmingly (∼70% of informative quartets) supports a (Scranciidae, Outgroup)|(Oenosandridae, (quadrifids, Notodontidae)) relationship. This quartet can be interpreted as topology A if rooted to the outgroups.

Incongruence is especially problematic for divergence time estimation (Carruthers et al. 2022), and we date three competing topologies to illustrate this problem. For example, the stem age of Scranciidae either antedates the K-T boundary or postdates it, depending on topology, which has major implications for the evolution of the group. In terms of biogeography, if Scranciidae are sister to all other Noctuoidea, this would suggest an Old World origin of the superfamily, but perhaps with little indication of more specific region considering the widespread extant range of Scranciidae. If, on the other hand, the Australian endemic Oenosandridae are the sister to the rest of the clade, as we found in some analyses, then the ancestral biogeography reconstruction may be even more restricted. Addressing the biogeographic origins of the most diverse Lepidoptera superfamily is obviously of interest but seems intractable with currently available data.

The transferal of *Scrancia* and its relatives from Notodontidae to their own family represents the discovery of a large, potentially independent lineage of trifids and carries significant ramifications for our understanding of noctuoid systematics and the characters on which it has traditionally rested. Regardless of the topology, trifid forewing venation can still be considered pleisiomorphic in Noctuoidea based on our results (Kitching and Rawlins 1998). But the well-established importance of the tympanal orientation, which has frequently been discussed in Noctuoidea systematics (Miller 1991; Kitching and Rawlins 1998; Zahiri et al. 2011), now falls into question, since the Scranciidae and Notodontidae share a ventral tympanal orientation, whereas quadrifids and Oenosandridae share the alternative backward facing state. This conundrum then requires different optimization of this trait since our data would suggest that the “notodontid orientation” could have evolved first in either the MRCA of Noctuoidea (if Scranciidae are sister to the remainder of the superfamily), or in the MRCA of Scranciidae + other clades (if Oenosandridae are sister to other noctuoids). Further highlighting the plasticity of this trait (and that of trifid venation), we note that Doidae were removed from Noctuoidea to the distantly related Drepanoidea, despite sharing the ventral-facing tympanal position of Scranciidae and Notodontidae, as well as trifid wing venation (Miller 1991; Kawahara et al. 2019; Rota et al. 2022). We include Doidae in our phylogenetic analyses (as an outgroup) as well as in Table S2 to stress the apparently homoplastic nature of these characters that have long been considered stable in Noctuoidea classification (e.g., Kitching and Rawlins 1998). Neuroanatomy of the tympana appears to be relevant to the systematics of Noctuoidea and would be an excellent direction of study for Scranciidae (Fullard 2006). Apart from its systematic utility, tympanal orientation also has far-reaching biological and evolutionary implications since the tympana are involved in detection and avoidance of bats and other predators (Kristensen 1998, 2003).

Scranciidae were discovered not by observing character conflict, which had never been conspicuous enough to question their placement in Notodontidae, but serendipitously, upon their inclusion in a phylogenomic analysis—a circumstance we anticipate repeating itself as more higher taxa are revealed by their inclusion in phylogenomic studies. Our hope is that the present work initiates a flurry of systematic, natural history, and evolutionary biology studies incorporating Scranciidae. Until now, little effort has focused on what was thought to be a rather obscure tribe, and then subfamily, of Notodontidae. Their putative placement as sister to the rest of Noctuoidea, wholly unique larval morphology, and geographic distribution all force a re-thinking of noctuoid evolution.

## SUPPLEMENTARY MATERIAL

Data available from the Dryad Digital Repository: http://dx.doi.org/XXX

## FUNDING

This work was supported by the Peter Buck Postdoctoral Fellowship at the Smithsonian Institution and SYNTHESYS+ (http://www.synthesys.info/) which is financed by European Community Research Infrastructure Action under the H2020 Integrating Activities Programme, Project number 823827.

## ACKNOWLEDGEMENTS

Ryan St Laurent thanks the Peter Buck Postdoctoral Fellowship and the SYNTHESYS+ for supporting this project. The following collections staff assisted with loans of material used in this study: Jim Hayden (MGCL) and Alberto Zilli (NHMUK). Specimens from Papua New Guinea and Nigeria were collected as part of other projects with the Binatang Research Centre and International Institute of Tropical Agriculture, respectively. *In situ* larval and adult images were provided by Vijay Anand Ismavel, Quartus Grobler, David Fischer, Wu Jiajie, SK Lai, Ian Sharp, Artur Tomaszek, Masatoshi Sone, and Cheryl Stinchcomb. Chris Owen (USDA) provided support with DiscoVista and genesortR. All authors would especially like to thank the late James Miller for his friendship and countless fruitful discussions about Notodontidae, not to mention his early recognition of scranciids as a unique higher-level taxon and exhaustive morphological study that paved the way for this work. Mention of trade names or commercial products in this publication is solely for the purpose of providing specific information and does not imply recommendation or endorsement by the USDA; USDA is an equal opportunity provider and employer.

## Notes

### Competing Interest Statement

The authors have declared no competing interest.

