## Supplementary material for "Hiding in Plain Sight: Phylogenomics Reveals a New Branch on the Noctuoidea Tree of Life": File S1

**File S1. Appendix 1.**

### SUPPLEMENTARY MATERIALS AND METHODS

#### *Morphology*

James Miller analyzed 100 adult and 74 larval characters of 50 notodontids (including two scranciids) comprising nine then-recognized subfamilies, two *incertae sedis* genera, and 13 outgroup exemplar noctuoids (Miller 1991). On that basis, Miller was the first to nominate a formal taxon for *Scrancia* and its relatives, the tribe Scranciini within the Dudusinae, and to establish criteria for testing its monophyly. We revisited Miller's matrix with a specific focus on the state coding and distributions of putative diagnostic characters of scranciids vis a vis Notodontidae and the other noctuoid families. Since our phylogenomic findings had indicated a placement of scranciids outside the Notodontidae, reexamining Miller's matrix served the trilateral purpose of (1) revisiting the putative synapomorphies of scranciids with Notodontidae; (2) specifying the impact of a novel topology on character transformations believed critical in noctuoid classification and evolution; and (3) refining the diagnosis of Scranciidae **stat. nov.** with re-interpreted character states and generating an enhanced framework for establishing the composition of the family.

We re-examined conflicts between putative synapomorphies of Notodontidae adduced by Miller (1991) and corresponding character states observed in scranciids, noting characters which, as originally coded, either (1) conflicted with inferred synapomorphies of Notodontidae or characters otherwise ubiquitous in Notodontidae; (2) agreed with those synapomorphies and therefore now potentially represent secondarily derived (homoplastic) characters whose reinterpretation as such is forced by our topological result; and/or (3) conflict with observation. Morphological data, including coding states are included in Table S2.

Genitalia preparations follow Clarke (1941) and Lafontaine (2004); some (not those figured) were stained with chlorazol black prior to slide-mounting in Euparal or maintained in glycerin. Dissections followed a brief (< 30-minute) heated soak in supersaturated NaOH. Wings that we examined were cleared by hand following soaking in a 10% bleach solution and then stained with eosin y. Wing preparation figures are derived from line drawings in Holland (1893a, 1893b) and Janse (1920), whereby Adobe Illustrator® in the Creative Cloud (Adobe Systems, Mountain View, CA) was used to trace the drawings and prepare vector illustrations.

We photographed genitalia with a Canon EOS 5Ds with an MP-E 65 mm lens on a stacking setup. The number of stacks were identified based on f-stop, lens, and magnification, adult specimens were photographed with the same camera and lens or a 100 mm lens for larger specimens. Images were edited with Adobe Photoshop® Creative Cloud and composed into plates using Adobe Illustrator® Creative Cloud. Slides and specimens used in morphological study are deposited at the USNM.

*Phylogenetics*

Partitions were first defined either by locus followed by codon position (60_percent_recovery_NA) or by locus only and consolidated further with ModelFinder. The preferred IQ-Tree run was selected out of 100 based on likelihood. For amino acid datasets, models were restricted to biologically relevant ones: Dayhoff, DCMut, JTT, JTTDCMut, LG, Poisson, PMB, VT, and WAG, but otherwise commands were the same as for nucleotide data.

*Dataset subsampling*

Subsampling of phylogenomic datasets is a common approach to reduce heterogeneity in the data and enhance reliability for phylogenetic inference (Smith et al. 2018; Mongiardino Koch 2021). So, as an alternative to selecting loci based on recovery, we also subsampled our data with genesortR, an R script that allows the user to sort hundreds of loci according to root-to-tip variance, saturation level, patristic distance, proportion of variable sites, average bootstrap, and Robinson-Foulds distance to a species tree (R Core Team 2020; Mongiardino Koch 2021). This method uses an input species tree (in this case the resultant ML tree from all_NA, locus partitioned analysis since all gene trees were analyzed), gene trees inferred with IQ-Tree unpartitioned and without SH-aLRT support values, and an alignment. Based on recommendations in genesortR, we set "topological_similarity = F" due to potential bias towards the provided relationships that may be uncertain, as in Noctuoidea; thus Robinson-Foulds distances are not accounted for in the subsampling procedure. Initial trials using genesortR with our data revealed a notable decline in locus quality (e.g. high root-to-tip variance, saturation, patristic distance, and relatively low values for proportion of variable sites and bootstrap) below the 600$^{th}$ ranked locus. Thus, our ninth and tenth (NSORT, AASORT) datasets are based on alignments of the 600 loci selected by genesortR, partitioned by locus with ModelFinder to merge partitions and identify best models of nucleotide evolution. Outlier locus removal was set to the default of 1% with outlier loci removed before the sorting step.

*Topology tests*

IQ-Tree topology tests calculate loglikelihood, delta loglikelihood (deltaL), bootstrap proportions Resampling of Estimated Log Likelihoods (bp-RELL), a p value for the Kishino-Hasegawa test (p-KH), a p value for the Shimodaira-Hasegawa test (p-SH), Expected Likelihood

Weight (c-ELW), and a p value for the Approximately Unbiased test (p-AU) as benchmarks. Since the KH test was designed for testing two trees only (Kishino and Hasegawa 1989) and the SH test is highly conservative when testing many trees (Shimodaira and Hasegawa 1999), we focus on the results of the AU test, which is meant to replace KH and SH tests as explained in IQ-Tree documentation.

*Divergence time estimation*

We do not utilize the Bayesian dating inference method in the study mentioned since it requires a fixed topology and greatly subsampled data. For the present study, the requisite subsampling (<100 loci) results in various poorly supported topologies (see main text discussions regarding our smallest datasets, for example). Given the high degree of uncertainty associated with node-dating and the inherently heterogeneous nature of target capture data, we adopt the less computationally intensive maximum likelihood (ML) method to date multiple topologies independently and examine the topological impact on estimated divergence times *a posteriori*. Following the pipeline in St Laurent et al. (2021), we first calculated 100 bootstrap replicates with RaXML (Stamatakis 2006) given the alignment and partition file from ModelFinder corresponding to the topology being dated. Once 100 bootstrap alignments and associated partition files were generated, these were used in all downstream analyses for each competing topology (a topology where Notodontidae are paraphyletic and the topology recovered only in parsimony analyses were omitted from dating). The second step in the pipeline requires input trees without branch lengths. For these, we use the previously discussed ML trees that gave rise to the various competing topologies being dated: Topology A–60_percent_recovery_NA (Scranciidae sister to remaining Noctuoidea); Topology B–

60_percent_recovery_AA (Scranciidae sister to [Notodontidae + Quadrifids]); Topology C–the 90_percent_recovery_NA (Scranciidae sister to Quadrifids). To remove branch lengths these trees were opened in FigTree v. 1.4.4 (Rambaut and Drummond 2009) and branch lengths set to "equal"; FigTree was used for all tree visualization as well. With branch lengths removed, we ran a priming TreePL step for 100 replicate bootstrap trees, followed by three independent cross-validation runs to identify the best smoothing parameter based on the chi-squared value; this was followed by a final complete TreePL run using the priming variable and smoothing parameters from the previous steps, resulting in 100 separate TreePL runs for each topology. TreeAnnotator 1.10.4 in the BEAST package (Drummond et al. 2012; Suchard et al. 2018) was used to combine each set of 100 dated trees.

For all dating analyses we used secondary calibrations from Kawahara et al. (2019), calibrating the same nine nodes as in St Laurent et al. (in press): The most recent common ancestor (MRCA) of Pyralidae and Notodontidae, 90.18–113.05 Ma; MRCA of Mimallonoidea, 18.63–45.7 Ma; MRCA of Mimallonoidea and Notodontidae, 86.4–108.37 Ma; MRCA of Drepanoidea, 76.84–98.94 Ma; MRCA of Bombycoidea and Lasiocampoidea, 74.15–94.4 Ma; MRCA Geometroidea, 72.31–93.08 Ma; the root age of Notodontidae (by calibrating using a notodontid and quadrifid), 66.97–88.57 Ma; an internal quadrifid node using *Helicoverpa zea* and *Manoba major* (Hampson), 53.7–73.82 Ma; and an internal branch considered the MRCA of Nystaleinae and Notodontinae (the two notodontid subfamilies present in Kawahara et al. (2019)), 31.52–66.07 Ma. The calibration for Drepanoidea was omitted for topology C since it was not recovered as monophyletic in the relevant ML run.

### SUPPLEMENTARY RESULTS

*Morphological corroboration*

Of all the Notodontidae subfamilies and tribes treated by Miller (1991), his Scranciini conformed least to the inferred synapomorphies of its parent family with the exception of the wing venation, tympanal, and stemapod characters, in part. Miller (1991: 185–186) wrote of scranciids that the adults “tend to be fairly small and light-bodied with elongate legs and wings. All larvae so far known have a large head, reduced prolegs on A3 and A4, and stemapodiform anal prolegs lacking crochets.” We note, first, that the reduced prolegs in combination with the presence of stemapods is unique among noctuoids to the Scranciidae with the exception of *Selenisa* (Hayward) and its relatives in the Omopterini (Erebidae: Erebinae). But in those taxa stemapods are likely independently evolved and are smoother with more truncated termini. In total, Miller (1991) presented eight adult and nine larval synapomorphies of “Scranciini”, but Miller’s diagnosis also applied to the subfamily diagnosis for Dudusinae (14 adult and 10 larval characters), in which he included the Scranciini. Therefore, in the main text we provide an updated diagnosis and a revised description restricted to true scranciids in the present appendix (below),

Recent works by Schintlmeister (2008) and Kobayashi and Nonaka (2016), establish the placement of Scranciids outside Dudusinae as we have. Our revised diagnosis of Scranciidae in the main text is therefore presented as an ensemble of Miller’s adduced synapomorphies of Scranciini and characters understood to be individually homoplastic but which in the aggregate are diagnostic of Scranciidae.

*Redescription of Scranciidae.*—Adults: *Head.* Ocelli large; male antennae pectinate to apex (Fig. S4e–g); female antennae ciliate; proboscis longer than thorax; R1 sensilla fluted; labial palpi moderate in length, segment 3 not unusually long; tentorium narrow, without crests;

frons protruding (Fig. S4e–g), strongly sclerotized, projections usually present. *Thorax.* Wings elongate, venation trifid, with a large, wide accessory cell (Fig. S4a–d). Legs long, foretarsi with first tarsomere longer than others combined; hind tibia approximately 1.5 times the length of the femur, spurs widely separated; tibial spurs in the formula 0-2-4; tarsal claws bifid. Female frenulum composed of fewer than 10 bristles. Metathoracic tympanum orientated ventrally, nodular sclerite absent. *Abdomen.* Anterior margin of tergum abdomen segment 1 with large internal bullae; cteniophores always absent from male abdomen segment 4; large internal bullae present on anterior margin of tergum of abdomen segment 1, dorsum of female abdomen segment 8 simple, pleuron sclerotized; posterior margin of male eighth sternite 8 often with a sclerotized notch; length of male tergite 8 equal to or shorter than that of tergite 7; terminal segments of male abdomen with a tuft of hairlike scales or a tuft of long pedicellate scales. *Male terminalia.* (Fig. S4i–l) Uncus/socii complex fused with tegumen, socii often absent; sacculus of male genitalia either lacking pleats or with faint pleats; ductus ejaculatorius simplex located posteriorly, with a long tubular projection formed by the anterior end of the phallus; cornuti absent from specimens examined. *Female terminalia.* (Fig. S4h) Ductus bursae usually sclerotized immediately below ostium, corpus bursae narrow usually with signum present, signum often coffee bean-shaped. *Larvae.* (Fig. 3) Head narrow, taller than thoracic segment 1 in lateral view (excluding legs), head wider than thorax in dorsal view; cranium narrow in lateral view, with a posterior depression along epicranial suture; head surface usually rugose and lacking secondary setae; labial palpus with a membranous flange on mesal margin; mandibular cutting edge smooth or serrate; spinneret dorsoventrally compressed, distal opening wide and flat; abdominal segment 3 prolegs smaller than those on abdominal segments 5 and 6, in some taxa nearly absent; abdominal segment 4 prolegs also smaller and with fewer crochets than prolegs of abdominal segment 5 and 6; abdominal segment 10 prolegs extremely long and flexible, stempodiform, crochets absent; proleg base with many setae, often with each on a raised tubercle; locomotion that of a looper (involving abdominal segments 4–6) or "semi-looper" (Fig. 3a, e) (involving abdominal segments 5–6; Holloway, 1983).

*Composition of Scranciidae*.—In the original description of Scranciini, Miller (1991) definitively included only *Gargetta* (Figs 1,2a, 3a–e, S4a) and *Scrancia* (Figs 1,2c,3h,i, S4a) because those were the genera examined as exemplars for character coding; some other genera were suggested to belong to Scranciini but not coded morphologically. Kobayashi and Nonaka's (2016) molecular phylogenetic study was the only work to include Scranciidae (as Scranciinae), and recovered them sister to Notodontidae based on up to two loci and with weak bootstrap support. These authors included *Gargetta*, *Hijracona* Holloway, 2011, *Lasioceros* Bethune-Baker, 1904 (Figs 1,2d,S4g), *Phycidopsis* Hampson, 1893 (Figs 1,2e), *Pseudogargetta* Bethune-Baker, and *Thacona* (Figs 1,2b) in their concept of Scranciinae (but did not sequence all of them), following Schintlmeister's (2008) elevation of the group to subfamily rank. Note that *Thacona* species were previously referred to as *Porsica* Walker throughout the literature, but *Porsica* was synonymized with *Thacona* in Holloway (2001). We agree with inclusion of the above genera in Scranciidae except for *Pseudogargetta* which we explain below. We also further broaden the concept of Scranciidae to include additional genera as mentioned in the main text (and Table S4) and remove others from the group.

A number of notodontid genera that resemble scranciids due to narrow wings have been formally treated as "Scranciinae" in the past, but do not belong to Scranciidae based on our molecular analyses and/or lack of observable Scranciidae apomorphies in those that we did not sequence: *Anticleapa* Kiriakoff, 1970, *Arceira* Kiriakoff, 1962, *Archigargetta* Kiriakoff, 1967,

*Baradesa* Moore, 1883, *Gargettiana* Gaede, 1930, *Ortholomia* Felder, 1861, *Polychoa* Turner, 1907, *Pseudogargetta* (Schintlmeister 2008, 2020; Schintlmeister and Witt 2015; Kobayashi and Nonaka 2016). We have included *Arciera*, *Pseudogargetta*, *Ortholomia*, and *Baradesa* in our phylogenetic reconstructions and each of these genera is recovered in notodontid subfamily Spataliinae (=Ceirinae of various authors, including Schintlmeister (2008), Schintlmeister and Witt (2015), and Miller et al. (2018)). *Ortholomia* and *Baradesa* have already been transferred to Spataliinae by Kobayashi and Nonaka (2016) and *Arciera* was transferred to "Ceirinae"(=Spataliinae) by Schintlmeister and Witt (2015). We therefore formally transfer *Pseudogargetta* to Spataliinae and putatively assign *Gargettiana*, *Polychoa*, *Archigargetta*, and *Antilceapa* to Spataliinae as well. These genera, while resembling Scranciidae superficially, can be differentiated by heavily sclerotized, well-developed socii, cteniophores in some (e.g. *Ortholomia*), stouter bodies, legs with the foretarsi no longer than the remaining tarsal segments combined, and absence of the large, broad, accessory cell on the forewing. The larva of *Archigargetta* is known and distinct from those of Scranciidae (Holloway 1983).

The following five genera may prove to belong to Scranciidae, but we have not been able to examine images or specimens sufficient to warrant formal assignment to any subfamily, so we treat them as *incertae sedis*: *Antiscranciola* Kiriakoff, 1967, *Mainiella* Kiriakoff, 1962, *Microchadisra* Kiriakoff, 1970, *Scaeopteryx* Kiriakoff, 1963, and *Scranciella* Kiriakoff, 1967.

Taxa from Asia and southern Africa have been revised (Schintlmeister 2008, 2020; Schintlmeister and Lourens 2010; Schintlmeister and Witt 2015), but Scranciidae of regions north of South Africa have not been studied systematically and many taxa remain undescribed. Although there is no complete taxonomic treatment on the included genera given above, there are roughly 100 valid species assigned to these genera in the global catalogue of Notodontidae (Schintlmeister 2013).

Holland W.J. 1893a. Descriptions of new species and genera of West African Lepidoptera. Psyche. 6:549–552.

Holland W.J. 1893b. Descriptions of new species and genera of West African Lepidoptera. Psyche. 6:531–538.

Holloway J.D. 1983. The Moths of Borneo: Family Notodontidae. Malay. Nat. J. 37:1–107.

Janse A.J.T. 1920. On the South African Notodontidae with descriptions of apparently new genera and species. Ann. Transvaal Mus. 7:149–237.

Kishino H., Hasegawa M. 1989. Evaluation of the maximum likelihood estimate of the evolutionary tree topologies from DNA sequence data, and the branching order in hominoidea. J. Mol. Evol. 29:170–179.

Kobayashi H., Nonaka M. 2016. Molecular phylogeny of the Notodontidae: Subfamilies inferred from 28S rRNA sequences (Lepidoptera, Noctuoidea, Notodontidae). Tinea. 23:1–83.

Lafontaine J.D. 2004. Noctuoidea: Noctuidae (part). Noctuinae (part — Agrotini). Fasc. 27.1. In: Dominick R., Ferguson D., Franclemont J., Hodges R., Munroe E., editors. The Moths of America North of Mexico, Fasc. 20.1. Washington, DC: Wedge Entomological Research Foundation. p. 385.

Mongiardino Koch N. 2021. Phylogenomic subsampling and the search for phylogenetically reliable loci. Mol. Biol. Evol. 38:4025–4038.

R Core Team. 2020. R: A Language and Environment for Statistical Computing. Vienna, Austria: R Foundation for Statistical Computing.

Rambaut A., Drummond A. 2009. FigTree v1.3.1.

Schintlmeister A. 2008. Palaearctic Macrolepidoptera 1: Notodontidae. Stenstrup: Apollo Books.

Schintlmeister A. 2013. World Catalogue of Insects, Volume II: Notodontidae & Oenosandridae (Lepidoptera). Leiden, Boston: Brill.

Schintlmeister A. 2020. The Notodontidae of the Indonesian Archipelago (Lepidoptera), Volume 1. Leiden, Boston: Brill.

Schintlmeister A., Lourens J.H. 2010. The Philippine Notodontidae (Lepidoptera). Quadrifina. 8:1–249.

Schintlmeister A., Witt T. 2015. The Notodontidae of South Africa including Swaziland and Lesotho (Lepidoptera: Notodontidae). Proceedings of the Museum Witt Munich. 2:1–289.

Shimodaira H., Hasegawa M. 1999. Multiple Comparisons of Log-Likelihoods with Applications to Phylogenetic Inference. Mol. Biol. Evol. 16:1114–1114.

Smith S.A., Brown J.W., Walker J.F. 2018. So many genes, so little time: A practical approach to divergence-time estimation in the genomic era. PLoS One. 13:1–18.

Stamatakis A. 2006. RAxML-VI-HPC: maximum likelihood-based phylogenetic analyses with thousands of taxa and mixed models. Bioinformatics. 22.

St Laurent R.A., Goldstein, P.Z., Miller J.S., Markee, A., Staude, H.S., Kawahara, A.Y., Miller, S.E., Robbins, R.K. Phylogenetic systematics, diversification, and biogeography of Cerurinae (Lepidoptera: Notodontidae) and a description of a new genus. Insect Systematics and Diversity. *In press.*

Suchard M.A., Lemey P., Baele G., Ayres D.L., Drummond A.J., Rambaut A. 2018. Bayesian phylogenetic and phylodynamic data integration using BEAST 1.10. Virus Evol. 4.

### Additional Supplemental Information

**Figure S1.** Schematic showing the seven competing topologies with regards to relative placement of Scranciidae, Oenosandridae, Notodontidae, and quadrifid families. The phylogenetic framework and dataset combination that give rise to each topology is/are listed beneath each topology.

**Figure S2.** Best of 100 IQ-Tree runs that gave rise to topology that was dated for Figure 1 in the main text. This tree shows the support values as UFBoot/SH-aLRT at all nodes. The tree file is available on Dryad.

**Figure S3.** The dated topology A, basis for Figure 1 in the main text, as output by a combined 100 TreePL runs summarized with TreeAnnotator. Mean ages are shown at all nodes, for full ranges of dates see the corresponding tree file on Dryad.

**Figure S4.** Relevant morphology of Scranciidae, a–d show wing venation (not to scale), e–f show heads, h–l show genitalia. All specimens in USNM (except those giving rise to the venation drawings). a) Male *Scrancia modesta* from Holland (1893). b) Female *Scrancia sticta* from Janse (1920). c) Male *Phycitimorpha stigmatica* from Janse (1920). d) Male *Dinotodonta longa* from Holland (1893). e) Male *Pseudoscrancia africana*. f) Male *Scrancia sp.* (antennae not shown). g) Male *Lasioceros aroa*. h) Female *Gargetta divisa*. i) Male *Gargetta breviosocius*, phallus below. j) Male *Pseudoscrancia africana*, phallus below. k) *Scrancia sp.*, phallus below. l) Male *Scrancia cf. astur*, phallus to left below that of Fig. S4k.

**Table S1.** Taxonomy, classification, and source of all samples used in the present study.

**Table S2.** Summary of character states bearing on the monophyly of Scranciidae and Notodontidae vis a vis other noctuoid clades: Oenosandridae, Doidae, and the quadirifids (Noctuidae, Erebidae, Nolidae, and Euteliidae). Character numbers follow Miller's (1991) treatment, except for those not treated by Miller (-), where 0/1 refer to absence/presence (other numbers are polymorphisms). Superscript "S" denotes synapomorphies of Scranciidae as enumerated by Miller (1991). Superscript "N" denotes synapomorphies of Notodontidae as enumerated by Miller (1991). An asterisk denotes predominant character state, if polymorphic.

**Table S3.** Summary of phylogenetic results, including data type, inference method (e.g. MSC, ML, or parsimony), phylogenetic program employed, dataset used, resultant topology, and all values of support for the roots of Scranciidae and Oenosandridae. In cases where a given topology was also dated (topologies A–C only) the crown and stem ages are reported for Noctuoidea, Scranciidae, Oenosandridae, quadrifid crown/stem age, and Notodontidae illustrating the impact of different topologies on divergence time estimates.

**Table S4.** Summary of genera hereby formally included in Scranciidae, those that may belong to Scranciidae, and those that are formally excluded and transferred to Spataliinae.
